# Continuity and discontinuity in human cortical development and change from embryonic stages to old age

**DOI:** 10.1101/329680

**Authors:** Anders M Fjell, Chi-Hua Chen, Donatas Sederevicius, Markus H Sneve, Håkon Gryde, Stine K Krogsrud, Inge Amlien, Lia Ferschmann, Hedda Ness, Line Folvik, Dani Beck, Athanasia M Mowinckel, Christian K Tamnes, René Westerhausen, Asta K. Håberg, Anders M Dale, Kristine B Walhovd

## Abstract

The human cerebral cortex is highly regionalized. We aimed to test whether principles of regionalization could be traced from embryonic development throughout the human lifespan. A data-driven fuzzy-clustering approach was used to identify regions of coordinated longitudinal development of cortical surface area (SA) and thickness (CT) over 1.5 years (n = 301, 4-12 years). First, the SA clusters were compared to patterns from embryonic cortical development. The earliest sign of cortical regionalization is the emergence of morphometric gradients in the cerebral vesicles, with a major gradient running along the anterior-posterior (AP) axis. We found that the principal divide for the developmental SA clusters extended from the inferior-posterior to the superior-anterior cortex, corresponding to the embryonic morphometric AP gradient. Second, embryonic factors showing a clear AP gradient were identified, and tests revealed significant differences in gene expression of these factors between the anterior and posterior clusters. Further, each identified developmental SA and CT cluster showed distinguishable lifespan trajectories in a larger longitudinal dataset (4-88 years, 1633 observations). This means that regions that developed together also changed together throughout life, demonstrating continuity in regionalization of cortical changes. The AP divide in SA development also characterized genetic patterning obtained in an adult twin sample, but otherwise regionalized CT development adhered more to the genetic boundaries. Finally, SA and CT clusters showed differential relationships to cognitive functions. In sum, the results suggest that development of cortical regionalization is a continuous process from the embryonic stage throughout human life.

**Significance statement (120 words):** The protomap and the radial unit theories of brain development have shown that graded expression patterns of several factors are responsible for shaping the ultimately highly partitioned and specialized neocortical landscape. The present study shows that the major anterior-posterior gradient of embryonic development can be detected in the regional cortical expansion profiles of children. The study further demonstrates that brain regions that develop together during childhood also tend to change together throughout the lifespan. This suggests that regional cortical development is a continuous process through the entire life, and that early-life factors have life-long impacts on this process.

---

Expansion of cortical surface area (SA) in human development is highly regionalized (1). According to the protomap (2) and radial unit models (3), the blueprint for this regional cortical expansion is established already in early embryonic development. Cortical neurons are not born within the cerebral cortex itself, but migrate from the ventricular zone to their final destination. The major morphometric gradient governing areal identity runs along the anterior-posterior axis (3). However, it is not known if the governing principles of embryonic brain development apply to cortical expansion in children. In the present study, we tested this hypothesis, which would show a continuity of cortical expansion from the early fetal to the latest stages of development in the pre-teen years.

Moreover, the related tenet that late-life brain health and cognitive function have developmental origins is getting increasing support (4, 5). Adult genetic SA topography - the delineation of regions influenced by the same genes - is characterized by an anterior-posterior gradient, likely due to prenatal factors (6, 7), corresponding to our hypothesis for organization of SA development. Thus, an intriguing possibility is that the fundamental organizational principles for regional brain development in children, likely reflecting patterns from embryonic development, can be traceable also in higher age. This would mean that anatomical regions that develop together also show distinct lifespan trajectories of adult cortical change and decline in aging. We know that cortical maturation to some degree proceeds along functional and structural networks established in adults (8−13), but not whether brain regions that develop together also change together through the rest of life. Testing this was the second main aim of the project.

These hypotheses were addressed through several steps. First, a data-driven fuzzy clustering approach was used to parcellate longitudinal change in SA and apparent thickness (CT) of the cerebral cortex into regions of coordinated development. CT and SA develop differently (1, 14, 15) and have different genetic and molecular foundations (2, 3, 16). We selected an age-range when SA still expands and apparent CT is continuously declining, i.e. 4-12 years (n = 301). Clustering was based on 1.5 years longitudinal change to avoid confounds from cross-sectional differences. The principal axes of the developmental clusters were extracted and tested against the anterior-posterior morphometric gradient from gene expression patterns in embryonic development, identified in (3).

The developmental clusters were further tested against genetic cortical topography established in an adult sample of twins (17, 18). CT shows more age change and is more affected by later-life events than SA (19−21), and the original genetic patterning work predicted that CT clusters, more than SA clusters, would show genetic relatedness with clusters of similar maturational timing (17). Thus, we hypothesized that developmental SA change would be relatively more influenced by prenatal factors while CT change clusters would be more similar to genetic clusters obtained from adult twins (22).

Next, we tested if the developmental clusters showed different age-trajectories in a longitudinal dataset covering more than eight decades (4.1-88.5 years, n = 974, 1633 scans). On the assumption of continuity between fetal and child development and lasting impacts of early-life factors on later brain changes, we hypothesized that the residual variance in each developmental cluster would be differentially related to age when the common variance shared among the clusters was accounted for. This means that we expected each developmental cluster to show different lifespan trajectories.

Finally, we tested whether the developmental clusters showed differential relationships to four empirically derived domains of cognitive function - episodic memory, executive-speed, working memory and general cognitive ability (GCA). These domains were identified from a principal component analysis (PCA) of multiple cognitive scores from an extended longitudinal dataset (4.1-93.4 years, 4065 observation). As SA and CT have different early determinants and show different developmental and lifespan trajectories, we hypothesized different relationships to cognition. We expected the SA clusters to relate to GCA, which is likely heavily influenced by early life factors and shows high life-span stability (5, 23−25). The scores included in the GCA component (matrix reasoning and vocabulary) load strongly on the g-factor (26), and have previously been found to correlate more with SA than CT (5, 23). As likely more amendable to environmental influences through life, CT was hypothesized to correlate more strongly with more specific cognitive functions, indicated by lower loadings on the g-factor. These could include episodic memory and executive-speed, which show different change trajectories across people (27) and are less strongly related to global cognitive change (28).

## Results

### Clusters of cortical change in development 4-12 years

Clusters are presented in Figure 1 (SA) and Figure 2 (CT). The two-cluster SA solution followed a clear anterior-posterior (AP, correlation with the AP axis r = .91) and inferior-superior (IS, r= .97) gradient, with less influence from the lateral-medial (LM, r = .19) axis. The AP division was tilted 19° from the base plane, causing the strong IS correlation. The posterior cluster covered the central sulcus and the adjacent anterior regions of the frontal cortex (premotor cortex), extending back to encapsulate the dorsolateral occipital cortex, and inferiorly down to the cingulum. For the three-cluster solution, an occipital-limbic cluster emerged. In the four-cluster solution, a new cluster appeared inferior to the first posterior cluster, distinguishing this cluster from the rest of the clusters both medially and laterally.

The two-cluster CT solution showed a very strong IS gradient (r = .97), with less AP (.39) and LM (.46) influence. With three clusters, an occipital-limbic cluster emerged. In the four-cluster solution, the superior cluster was split in two clusters. For both CT and SA, the silhouette values flattened after 4 clusters, suggesting that 5 clusters were too many (see SI). The adjusted Rand Index (adj RI) showed substantial overlap between the 2-cluster SA and CT solutions (adj RI = .88), with less similarity for three (adj RI = .27) and four (adj RI = .22) clusters. Still, the medial limbic cluster and the prefrontal - anterior-temporal relationship was similar between SA and for CT across the 3- and 4-cluster solutions.

**Figure 1.**
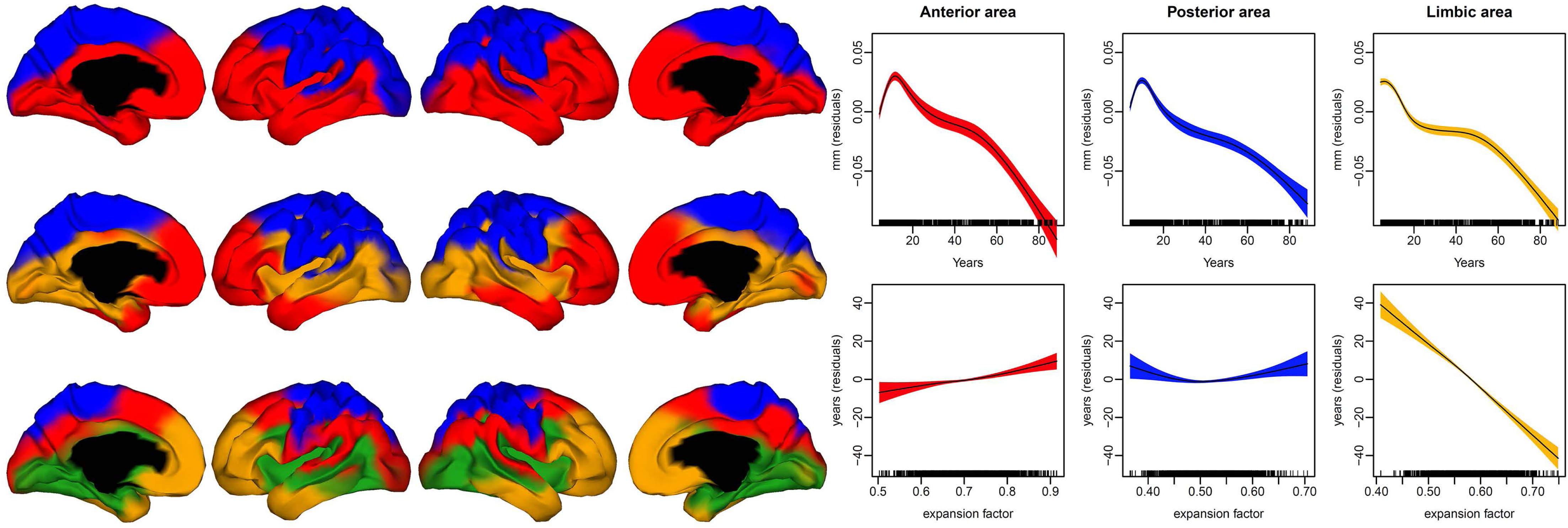
Surface area. Left panel: Clusters of coordinated surface area in development, two-(top), three-(middle), and four-(bottom) cluster solutions. Right panel: The lifespan trajectories of each cluster from the three-cluster solution. Top: Trajectories residualized on age (x-axis). Bottom: The residual age-relationship (y-axis) for each cluster accounting for the other two clusters. The colors of the curves correspond to the cluster color in the left figure. The shaded area denotes +/-2 standard errors of the mean.

**Figure 2.**
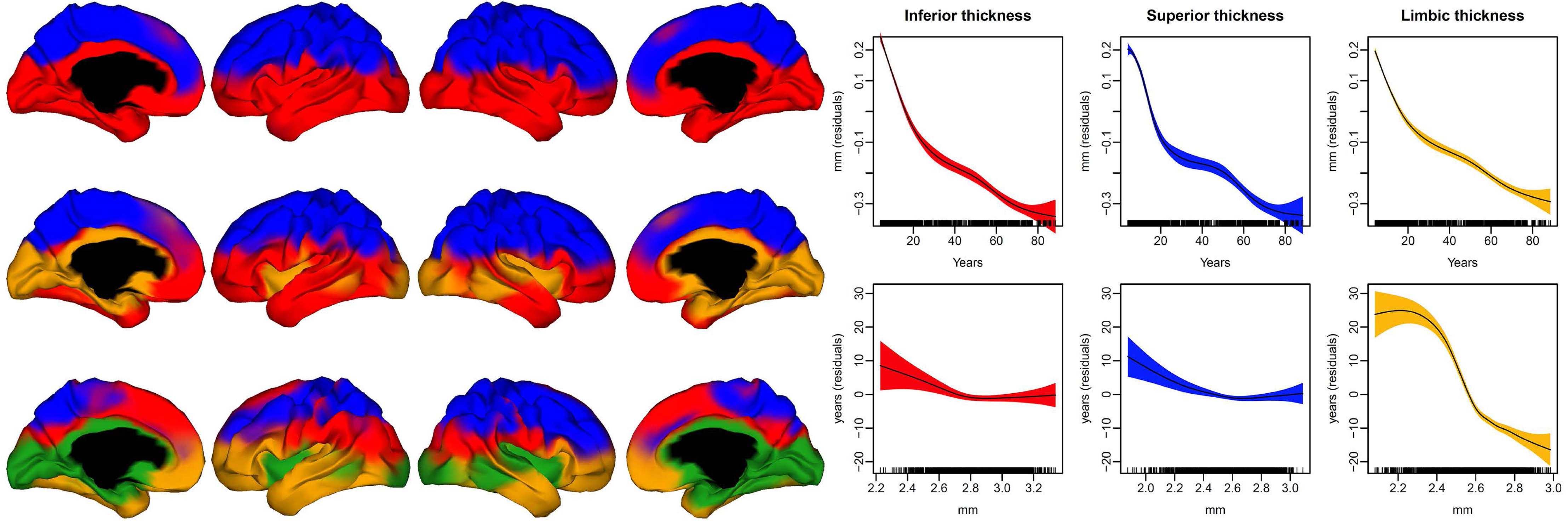
Cortical thickness. Left panel: Clusters of coordinated cortical thickness in development, two-(top), three-(middle), and four-(bottom) cluster solutions. Right panel: The lifespan trajectories of each cluster from the three-cluster solution. Top: Trajectories residualized on age (x-axis). Bottom: The residual age-relationship (y-axis) for each cluster accounting for the other two clusters. The colors of the curves correspond to the cluster color in the left figure. The shaded area denotes +/-2 standard errors of the mean.

To validate the Fuzzy clustering results, we ran seed point analyses from 360 seeds based on a multimodal parcellation scheme (29). The critical features of the cluster results were confirmed, including the AP axis for SA, the IS axis for CT, and for both the prefrontal-temporal relationship and delineation of the limbic clusters (see SI).

### Adherence of SA development to early morphometric gradients

We rated 31 transcription factors assumed responsible for shaping the partitioned neocortex (3) according to how well their expression patterns were aligned to the AP axis. 18 factors showed a clear and four a partial AP gradient, yielding evidence that the AP axis constitutes a major regionalization gradient in early cortical development, resembling the AP axis of the SA clusters in later child development. To test whether this cluster gradient could be detected in human gene expression patterns, we selected the candidate with the most established anterior-posterior expression pattern, PAX6 (30), and the candidate with the most pronounced posterior-anterior expression pattern, P75 (3). Mean expression values for cortical regions falling within the anterior and the posterior developmental clusters were extracted for 34 participants from the Brainspan Atlas (www.brainspan.org. see SI). Expression values of PAX6 were significantly higher (t = 4.24, df = 33, p < .0002) in the anterior than the posterior development cluster, while the opposite was found for P75 (t = −3.03, df = 33, p < .005) (Figure 3). Put together, this shows that traces of the major expression patterns in embryonic development can be recovered in later childhood, and that the gene expression patterns in humans follow cortical change clusters in ways predicted from early embryonic development.

**Figure 3.**
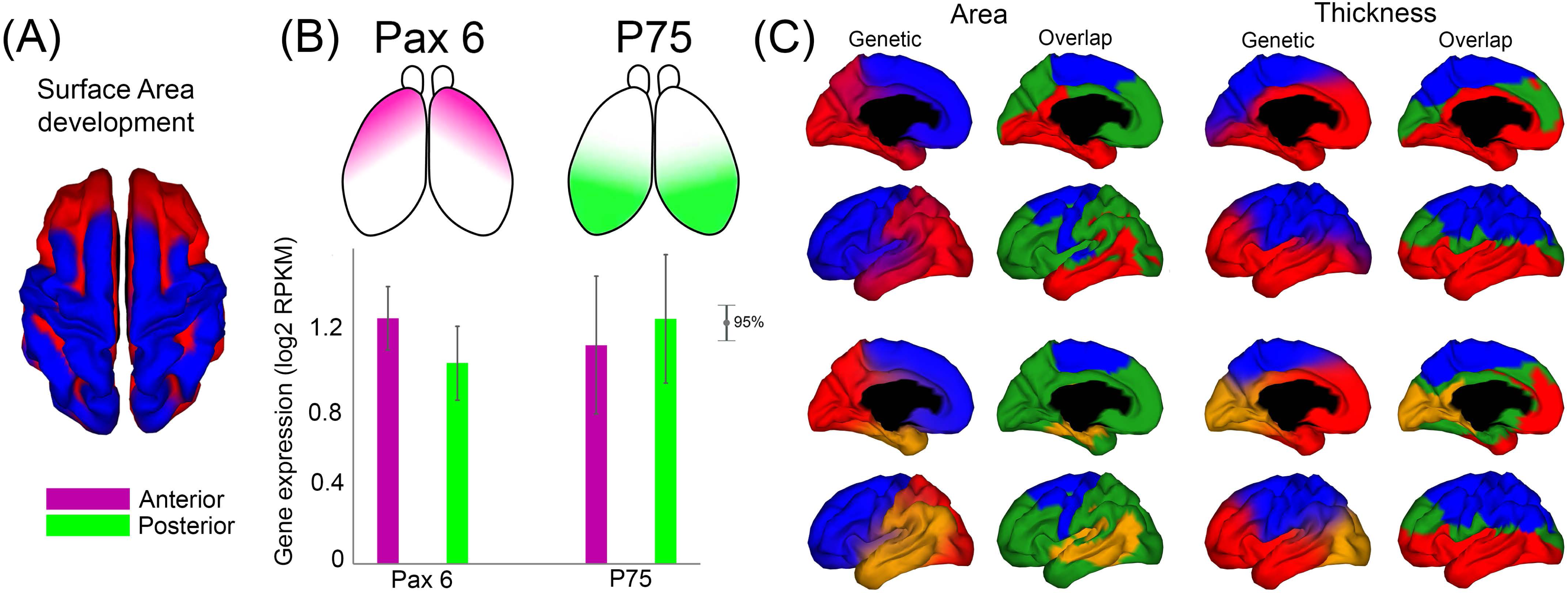
Genetic patterns. A: Surface area (SA) developmental clusters, top view. B: Anterior-posterior gradient of PAX6 (left) and P75 (right) from (3). The error bar plot (bottom) shows higher expression of Pax6 in the anterior cluster and higher expression of P75 in the posterior cluster in the Lifespan database. C: Genetic clusters from (17) for SA (left) and thickness (right), two-(top) and three-(bottom) solutions. The overlap plots show vertices with the same cluster identity across genetics and development in blue/ red/ yellow, and the vertices with different identity in green.

### Resemblance to cortical genetic patterning

After having established that SA development follows the main AP gradient identified in embryonic development, we tested how similar the developmental SA and CT clusters were to mid-life genetic cortical topography (17) (Figure 3). Genetic topography represents the delineation of regions influenced by the same genes. Rand Index suggested low overlap (adj RI < .10 for 2 and 3 cluster solutions) between genetic topography and the developmental clusters. The AP division was seen for both, but the boundary between the clusters was shifted from inferior-posterior to anterior-superior for the development clusters (−71° from the perpendicular axis) compared to inferior-anterior to superior-posterior (+38°) for the genetic clusters. Adherence to the AP axis was slightly higher for the genetic (r = .96) vs. the developmental (r = .91) clusters, and slightly lower for the IS axis (r = .82 for genetics vs. r = .91 for development).

Comparing the developmental and the genetic CT clusters, the same IS division was seen for the 2-cluster solution (r = .96 for genetics vs. .93 for development, RI = .69, adj RI = .39). The genetic clusters also followed the AP (r = .90) and to a lesser extent the medial-lateral (r = .25) gradient. For the 3-cluster solution, the medial delineation of the limbic developmental cluster included the medial temporal lobe and the cingulum as well as the occipital cortex, while the genetic homologue cluster did not include the occipital lobe but more of the prefrontal cortex laterally and medially. RI indicated similar genetic-developmental overlap (RI = .68) for the three-cluster solutions, but adj RI dropped to .31, with a further drop to the 4-clusters solution (RI = .70, adj RI = .23). Thus, the overlap between CT development and genetic topography dropped with higher number of clusters.

### Lifespan-trajectories of developmental clusters

Mean CT and SA values in each cluster from the 3-cluster solutions were extracted for each participant and time point in the full MRI sample from 4.1 to 88.5 years, yielding 1633 observations. Generalized Additive Mixed Models (GAMM) with cluster as dependent variable, age as predictor and a random effect for intercept, showed that all clusters showed highly non-linear relationships to age (all p’s < 2e^−16^, effective degrees of freedom [edf] for all ≥ 5). Sex impacts SA (1), and was included as a nuisance covariate in the SA analyses. As expected, CT clusters showed mainly monotonous negative relationships to age (Figure 2), with steeper reductions in childhood and adolescence, while SA increased in the first part of the age-span (Figure 1), peaking in early teenage years, followed by reductions for the rest of the life-span.

To test whether the clusters showed unique age-trajectories, analyses were repeated with all clusters for each modality (SA, CT) as simultaneous predictors of age. For SA, this yielded highly significant residual age-relationships for all clusters (all p’s < 4.07^−5^). The limbic cluster showed residual reductions with higher age throughout the whole age-span, while the anterior cluster showed slight increases and the posterior a U-shaped trajectory (Figure 1, right panel, bottom). The analyses were also re-run without the cluster-forming development sample, and all clusters were still highly significantly related to age when run separately (all p’s < 2e^−16^) and simultaneously (n = 502, 860 observations, all p’s < .05), both for CT and SA.

### Random clusters

We created random clusters of approximately the same size and repeated the GAMMs with all clusters as simultaneous predictors of age (see SI). The developmental clusters performed substantially better than the random clusters on all model fit measures (CT: developmental clusters - random clusters, ΔAIC = −150.75, ΔBIC = −150.75, logLik = 75.38/ SA: ΔAIC = −53.57, ΔBIC = −53.58, logLik = 26.79).

In conclusion, we were able to detect large cortical regions with unique and independent trajectories across the lifespan through data-driven clustering of longitudinal change in children. We proceeded to test whether the clusters were related to cognitive performance.

### Cognitive components - relationship to developmental clusters

All cognitive components showed highly significant (p < 2e^−16^) and non-linear (edf ≥ 4.9) age-trajectories (Figure 4A, see SI). Each component was then used as dependent variable in separate GAMMs, with age and each developmental cluster as predictors, intercept as a random effect and subject time point as covariate to control for retest effects. Selected results are presented in Figure 4C (full results in SI). All SA clusters were positively related to GCA (p < .0002). The anterior cluster was also related to working memory span (p = .008), while the limbic (p = .03) and posterior (p = .018) did not survive Bonferroni corrections. None were related to memory or executive-speed. In contrast, the CT superior cluster (p = .005) was related to memory, while the limbic and inferior (both p’s = .02) did not survive corrections. The limbic (p = .01) and inferior (p = .005) were related to executive-speed (after corrections). The relationships were non-linear, as would be expected based on the monotone negative thickness-age relationships coupled with the typical inverse U-shaped cognition-age relationships. The differential relationships between cognitive function and CT and SA are illustrated in the heat maps in Figure 4B.

**Figure 4.**
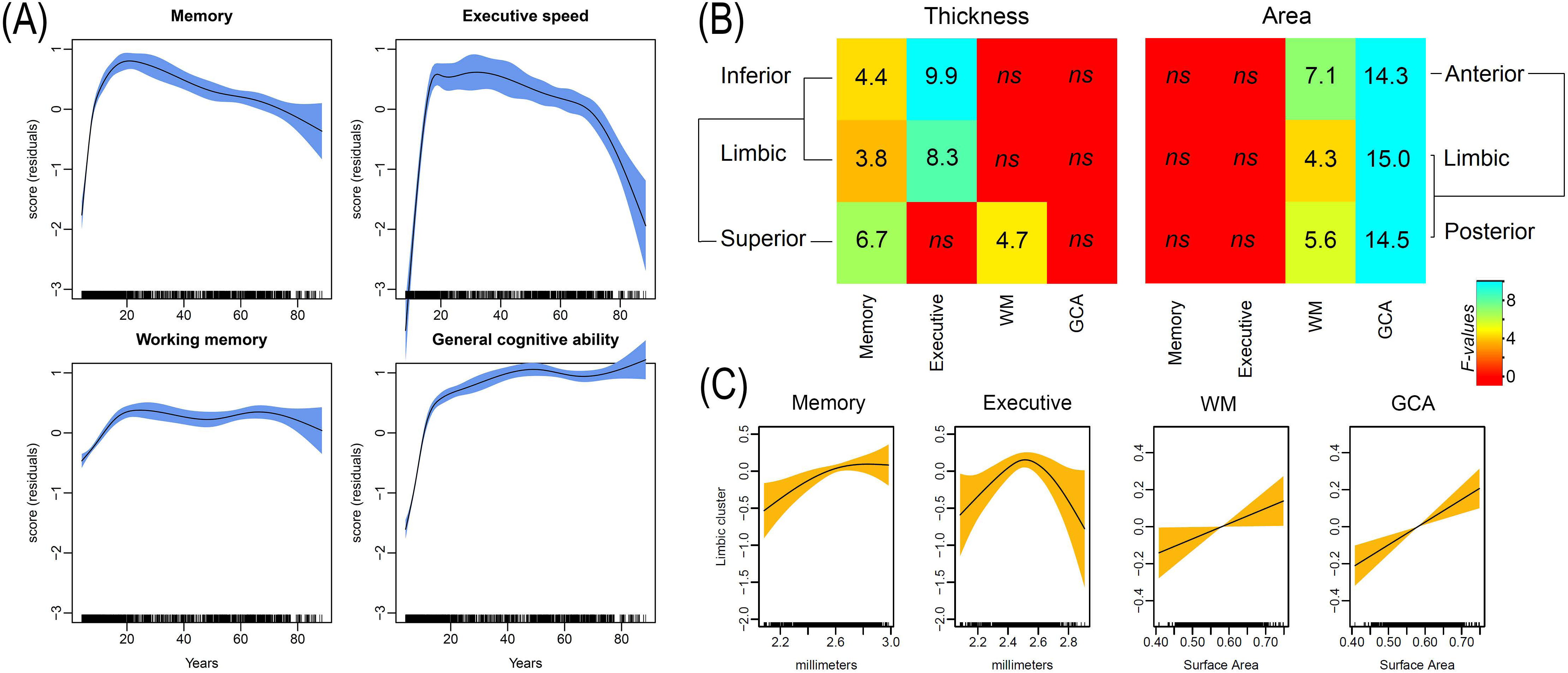
Cognitive relationships. A: Each cognitive domain plotted against age, residualized on time point to remove retest effects. For trajectories of the tests loading highly on GCA (matrix reasoning, vocabulary) see SI. B: Heat maps of F-values illustrating the relationship between each cluster and each cognitive function, regressed on age and sex (SA). C: Examples of brain cognition-relationships for the limbic cluster for thickness (two plots to the left) and area (two plots to the right). Plots for all variables are presented in SI. The shaded area denotes +/-2 standard errors of the mean.

## Discussion

The present results show that cortical expansion in childhood development follows the major gene expression patterns of factors responsible for shaping the partitioned and specialized human neocortex (3). Through a data-driven parcellation of the cerebral cortex based on longitudinal change in SA in 4-12 year old children, we were able to detect the major anterior-posterior developmental gradient known from embryonic development (30). This demonstrates a continuity in cortical expansion from the earliest stages to the period when regional arealization reaches its peak. While this represents continuity between the fundamental principles for embryonic brain development (2, 3) and later childhood SA maturation, in line with known influences of neonatal characteristics on SA (7), CT change more strongly followed adult genetic organization principles, as predicted from previous work on genetic cortical patterning (17). Importantly, the developmental clusters showed statistically distinguishable trajectories through more than eight decades of life, in accordance with the view that fundamental principles governing brain development in children have lasting impact on the brain. Thus, brain regions that develop together also change together through the rest of life. Finally, since SA is highly determined by prenatal factors, we expected correlations with GCA, which loads highly on the g-factor (26) and has high between-person lifespan stability (24, 25). This was confirmed, in contrast to CT, which was more strongly related to cognitive domains with lower loading on the g-factor (26), such as memory and executive-speed. This suggest that cortical characteristics (arealization and thickness) beyond anatomical boundaries are the more important in defining relationships to cognition.

### Parcellation of the developing human cerebral cortex

The results showed that the cortex can be parcellated in meaningful ways based on developmental change data alone. The anterior-posterior SA axis followed the major gene expression gradient identified in animal studies (3, 30). This link was further supported in the present study through the finding of significantly different gene expression in the anterior vs. the posterior cluster for PAX6 and P75, both established anterior gradient factors in embryonic development (30). Such correspondence between childhood SA maturation and cortical ontogenesis is in line with the protomap (2) and radial unit tenets (3). According to these, neural progenitors in the ventricular zone form a mosaic of proliferative units that establish an embryonic cortical map in early development. During migration of neurons into the cortex, their position is then retained by restraints imposed through the radial glial scaffolding. However, it has not been established that these principles laid down in the first stages of CNS development are reflected in cortical maturation in later childhood in humans.

Although there was good overlap between SA and CT for the two-cluster solution, this dropped substantially with higher cluster numbers. CT and SA are shaped by independent genes (2, 16) and different neurobiological mechanisms in early fetal life (3), and they also follow fundamentally different trajectories in development (1, 14, 15) and aging (20). As expected, there was a fundamental difference in the principal axis distinguishing CT and SA even for the two-cluster solution, with the CT clusters adhering to an inferior-superior axis, with the plane of the cluster division tilted 19° for SA. While the anterior-posterior SA division was seen also in a previous developmental study, the inferior-superior CT axis was not (10). This could possibly result from methodological differences, e.g. that the previous study used both combined cross-sectional and longitudinal information. A second longitudinal study with an alternative analysis approach reported indirect evidence for an inferior-superior gradient in CT development (13), and it has previously been suggested that such a division could be due to cytoarchitectonic features and connectivity patterns (17).

The three- and four cluster solutions revealed additional interesting patterns. Not only local clusters were seen, but also relationships between functionally connected regions crossing lobar boundaries. For instance, lateral prefrontal cortex and the lateral anterior temporal cortex were included in the same SA and CT clusters. This fronto-temporal connection fits earlier observations (8, 10), and would be expected from structural connections represented by the uncinate fasciculus (31). A limbic cluster encapsulating the medial temporal lobe, the cingulum and insula, and extending backward covering the lingual gyrus, cuneus and occipital cortex, was also seen for both SA and CT. This demonstrates that cortical maturation not necessarily follows lobar boundaries, but rather extends beyond what would be expected from anatomical proximity. Similarly, SA of auditory cortex in the superior temporal lobe clustered together with somatosensory, motor and visual cortices in the two-cluster solution, i.e. mimicking functional clusters. Still, morphological change did not follow functional boundaries in a clear-cut manner, as calcarine sulcus (V1) and ventral visual areas clustered with the inferior/ anterior cortex, not the rest of the visual cortex.

Importantly, the developmentally defined clusters showed statistically distinct lifespan trajectories. Although the general form of the trajectories is well known (5, 22), the analyses revealed residual age-relationships for single clusters independently of other clusters. This finding was in accordance with the hypothesis that variations in cortical development has life-long impacts, and consequently that regions that develop together tend to change together through the rest of life. We have previously shown that correlations between longitudinal changes in CT across predefined smaller ROIs are similar in development and aging (22). Here we show that the trajectories of clusters based solely on coordinated developmental change, with no anatomical restrictions imposed on the clustering, can be delineated in a sample spanning more than 80 years, both for SA and CT.

### Adherence to genetic patterning

The clusters were tested against genetic cortical patterning results (17). For CT, we found overlap between developmental and genetic patterns. This fits with our previous finding that regional differences in cortical thinning adhered to genetic organizational principles (22). In the present study, we defined clusters based on developmental change, independently of the genetic patterning. Although the genetic overlap was lower with a higher number of clusters, certain similarities between developmental and genetic organization patterns remained, for instance the prefrontal-temporal relationship (17). Thus, partly the same genes seem to govern absolute frontal and temporal CT. A genetic fronto-temporal relationship was also seen in a developmental twin sample for CT and SA (32).

In contrast, there was limited overlap between the developmental SA clusters and the genetic clusters, revealing interesting differences. Genetic SA relationships seem local or lobar, lacking strong long-distance or cross-region correlation patterns, with generally lower - and more age-invariant - correlations across the cortex (32−34). The developmental organizational patterns crossed anatomical boundaries, such as the fronto-temporal and the limbic clusters. Another difference was that premotor and postcentral/ somatosensory regions clustered together in development, as seen in previous fetal work (35), but have distinct adult genetic patterns (17). Notably, the genetic-developmental similarity for CT is in line with the prediction from the original genetic patterning paper that CT clusters would show genetic relatedness with clusters of similar maturational timing (17).

The difference in genetic overlap for CT and SA further illuminates the developmental origins hypothesis. We expected SA change to adhere to major gradients from embryonic development, and CT change to overlap with genes responsible for CT at a later point in life. SA reaches its maximum expansion in early adolescence (1), with the major determinant likely being the number of cell division cycles of the neural stem cell pool at the premitotic stage of neural development (6). The width of the cortical mini-column at birth is around one-third of its adult size (36), and expansion of cortical SA in childhood can likely be explained by growth of these (37). The developmental SA clusters likely reflect the last part of an expansion that can be traced back to embryonic stage, as also evidenced by the demonstrated adherence to the anterior-posterior gradient of gene expression patterns. In contrast, CT ceases to increase early after birth and reaches 97% of adult values in two years (37). Early CT is affected by asymmetric cell division cycles of the cortical founder pool, determining the number of cells within each column (6). However, the number of neurons in each column is hardly a determinant of cortical changes after the initial CT increase, which rather can be attributed to dendritic branching, synaptogenesis and gliogenesis (38). Thus, CT change from 4 to 12 years likely reflects either different processes or differences in the relative contributions of the same processes compared to CT change in very early development. Studies show that the genetic contribution to CT but not SA changes during development (32, 39), and CT shows steeper slopes during adolescent development (1) and later aging (20) than SA. We argue that CT to a greater extent than SA reflects accumulated life-long genetic and environmental impact. Thus, SA is usually more strongly related to early life factors and CT relatively more also to later life factors. Empirical support comes from studies showing larger effects of birth weight and other obstetric factors on SA than CT (7, 40−42), and environmental interventions affecting CT (19, 21).

### Cognitive correlates of developmental clusters

SA and CT were expected to correlate with different cognitive domains. We hypothesized that SA would be related to GCA, which shows high between-person stability across life (24, 25), likely due to the impact from early life-factors. Accordingly, we found highly significant lifespan relationships between GCA and the SA clusters, but not CT clusters, in line with previous observations (5, 23). In contrast, we speculated that since CT is more strongly related to age and sensitive to environmental influences, it would be related to more specific cognitive functions with lower loadings on the *g-* factor and less established lifelong between-person stability. Episodic memory and executive-speed have lower g-loadings than GCA (26) and declines steeply with age (43). We found relationships between these components and all CT clusters. No significant relationships were seen between executive-speed and SA, while the relationships between SA and memory were just significant (limbic and posterior clusters) or showed a trend (p = .06, anterior cluster). Working memory span, which was relatively invariant during adult life, showed comparable relationships with CT and SA. Importantly, however, greater age effects do not equal less between-person stability, and it is as of yet not clear whether memory and executive-speed show less between-person stability over the life-course than GCA. People differ in the amount of change in memory and speeded tasks over time (27), and change in GCA is more strongly related to “global cognitive” change than changes in episodic memory and speed are (28). Still, correlations of change within and across cognitive domains tend to increase with age (44), and a recent meta-analysis showed that 59% of the variance in change is shared across abilities (45). In any case, a conclusion from the present results is that relationships between cognitive function and cortical morphometry adhere more strongly to modality than anatomical region. Vertex-wise analyses across the cortex could potentially detect more anatomically specific relationships with cognition, but at the levels of major clusters, measurement type seems more important than anatomical location.

### Conclusion

In conclusion, cortical SA expansion during childhood was organized according to an AP gradient, seen also during early embryonic development, and related to gene expression and genetic patterning. Clusters of cortical development showed statistically distinct trajectories through 8 decades of life, and correlated with cognitive function in predictable ways, demonstrating continuity of human cortical development from early to late stages of life.

## Materials and methods

### Sample

A total of 1633 valid scans from 974 healthy participants (508 females/ 466 males), 4.1 to 88.5 years (mean visit age 25.8, SD 24.1), were drawn from three Norwegian studies coordinated by the Research Group for Lifespan Changes in Brain and Cognition; The Norwegian Mother and Child Cohort Neurocognitive Study (MoBa)/ Neurocognitive Development (ND)/ Cognition and Plasticity Through the Lifespan (CPLS)) (see SI for details). 635 participants had two scans and 24 had 3 (mean scan interval 2.3 years [0.2-6.6]). The sample is identical to (22). All were screened for conditions assumed to affect CNS function. The cluster-forming sample consisted of all MoBa participants with two scans (n = 301, age 7.3 [4.1-12.0], mean scan interval 1.5 years [1.0-2.2]). The twin sample used to generate the genetic clusters consisted of 406 middle-aged men (55.8 years [51-59]), including 110 monozygotic and 93 dizygotic twin pairs, from the Vietnam Twin Study of Aging (see SI and (46) for details).

### MRI data acquisition and analysis

Imaging data (except VETSA) were acquired using a 12-channel head coil on two 1.5-Tesla Siemens Avanto scanners (Siemens Medical Solutions, Erlangen, Germany), yielding 2 repeated 3D T1-weighted magnetization prepared rapid gradient echo (MPRAGE): TR/TE/TI = 2400 ms/ 3.61 ms/ 1000 ms, FA = 8°, acquisition matrix 192 × 192, FOV = 192,160 sagittal slices with voxel sizes 1.25 × 1.25 × 1.2 mm. For most children 4-9 years old, iPAT was used, acquiring multiple T1 scans within a short scan time. For VETSA, images were acquired on two Siemens 1.5 Tesla scanners. Sagittal T1-weighted MPRAGE sequences were employed with a TI=1000ms, TE=3.31ms, TR=2730ms, flip angle=7 degrees, slice thickness=1.33mm, voxel size 1.3×1.0×1.3mm. MRI data were processed and analyzed with FreeSurfer 5.3 (http://surfer.nmr.mgh.harvard.edu/) (47, 48) longitudinal stream (49). SA maps were smoothed using a circularly symmetric Gaussian kernel with a full width at half maximum (FWHM) of 26 mm and CT maps with a kernel of 21 FWHM. See SI for details. Movement is a major concern (73), so scans were manually rated 1-4, and only 1 or 2 (no visible or only very minor possible signs of movement) were included.

### Fuzzy clustering

The fuzzy clustering procedure was performed using the ‘cluster’ package implemented in R (www.rproject.org/). The individual de-meaned annualized symmetrized percentage change (APC) maps were fed into the cluster algorithm. We calculated pair-wise correlations of CT and SA change between every two vertices on the cortex for the left and right hemispheres separately.

### Cognitive testing

Principal component analysis (PCA) was run on the cognitive test scores from a larger sample from the same research center (n = 4065 observations, inclusive of the 1633 MRI subsample observations, age 4.1-93.4 years, 2285/ 1722 females/ males observations) to extract four stable general domains of cognitive functions, yielding 81.53% explained variance. Procrustes rotation was used to confirm that the PCA was representative also for the MRI subsample (correlation of .95, p < 9.999e^−05^). See SI for details.

### Statistical analyses

Analyses were run in R (https://www.r-proiect.org) using Rstudio www.rstudio.com) IDE, except the PCA (SPSS v25). The adjusted Rand index (RI) (50) and Silhouette plots, in combination with visual inspections, were used to inform the choice of specific cluster solutions. CT and SA data from each cluster were extracted for all participants in the full sample and were averaged across hemispheres. GAMMs using the package “mgcv” (51) were used to derive age-functions. Akaike Information Criterion (AIC) (52) and the Bayesian Information Criterion (BIC) was used to guide model selection. See SI for details.

## Acknowledgement

This work was supported by the Department of Psychology, University of Oslo (to K.B.W., A.M.F.), the Norwegian Research Council (to K.B.W., A.M.F.) and the project has received funding from the European Research Council’s Starting Grant scheme under grant agreements 283634, 725025 (to A.M.F.) and 313440 (to K.B.W.). We thank Jon Skranes for organizing the NTNU part ofMoBa.

